# Test-retest reliability of perturbation-evoked cortical activity reflects stable individual differences in reactive balance control

**DOI:** 10.1101/2024.10.03.616575

**Authors:** Jasmine L. Mirdamadi, Alex Poorman, Gaetan Munter, Kendra Jones, Lena H. Ting, Michael R. Borich, Aiden M. Payne

## Abstract

There is a growing interest in measuring cortical activity during balance control for understanding mechanisms of impaired balance with aging and neurological dysfunction. The most well-characterized electrophysiological signal elicited by a balance disturbance is the perturbation-evoked N1 potential that peaks 100-200 msec post-perturbation and is thought to reflect error processing. We previously found associations between the N1 and individual differences in balance ability, suggesting it may be a potential biomarker of balance health. However, a potential biomarker of balance function will be limited by its reliability and clinical feasibility, which has yet to be established. Here, we characterized the reliability of the N1 elicited by standing balance perturbations within and between sessions over a one-week interval in 10 younger and 14 older adults. A subset of older adults (n=12) completed a session approximately one year prior to the main experiment. We extracted N1 amplitude and latency from the Cz electrode using an advanced, computationally-intensive approach that relies on large amounts of data (e.g., 64 channels, many trials), Test-retest reliability was assessed using the intra-class correlation coefficient (ICC). Internal consistency was quantified by split-half reliability using the Spearman correlation coefficient. N1s varied across individuals (amplitude:12-82µV, latency: 142-282ms), yet within individuals, the N1 showed excellent test-retest reliability (ICC>0.9) across a one-week and one-year time span. N1 amplitude reached excellent internal reliability for each session and group (r>0.9), that generally plateaued within 6 trials, while more trials were needed to reliably measure N1 latency. Similar results were obtained when quantifying N1s using a minimal approach performed with only three electrodes and simple preprocessing. Overall, the N1 is stable within and across sessions, and is largely independent of approach suggesting it could be a clinically-feasible biomarker of balance function. Characterizing reliability in populations with neurologic dysfunction and in different environmental contexts will be necessary to enhance our understanding of the N1, optimize experimental design, and determine its predictive validity of clinical outcomes (e.g., falls risk).

## 1. Introduction

Individual differences in brain activity during balance recovery behavior may reveal brain mechanisms of balance impairment and fall risk with aging and neurological conditions. Electroencephalography (EEG) can quantify temporally-precise patterns of cortical activity on physiologically-relevant time scales during balance recovery behavior. The most well-characterized EEG signal elicited by a sudden balance disturbance is the perturbation-evoked N1 potential, a negative voltage deflection that typically peaks 100-200 msec post-perturbation (Purohit and Bhatt, 2022; Varghese et al., 2017). The N1 is thought to reflect a sensory-driven error assessment signal that can be elicited in different contexts (sitting (Mochizuki et al., 2009), walking (Quintern et al., 1985), lean-and-release (Mochizuki et al., 2010), floor tilt, support-surface translations (Payne et al., 2019a)) and in different populations (toddlers (Berger et al., 1987), younger adults, older adults (Payne et al., 2023, 2021), Parkinson’s disease (Payne et al., 2022), stroke (Palmer et al., 2024). Several studies have explored factors that modulate the N1 within individuals, including predictability (Adkin et al., 2006; Mierau et al., 2015), task difficulty (Payne and Ting, 2020; Solis-Escalante et al., 2021), perceived threat (Adkin et al., 2008), and attention (Quant et al., 2004). However, such within-subjects investigations cannot establish whether the N1 is suitable for distinguishing between individuals, which is necessary for most clinical applications. More recently, between-subjects investigations have linked the N1 to individual differences in balance (Payne and Ting, 2020) and cognitive (Payne et al., 2021) abilities, but have yet to assess the reliability of the N1 as a stable measure of individual differences in reactive balance control, a necessary step towards optimizing experimental designs in basic science and for advancing its potential clinical utility as an electrophysiological biomarker of brain function related to balance health.

For a brain measure to be clinically useful, it must be relatively stable over clinically relevant timescales, which can be assessed in terms of test-retest reliability (McEvoy et al., 2000). Changes in brain activity may precede behavioral declines (Beason-Held et al., 2013) and be more sensitive to subtle changes in neural mechanisms underlying behavioral changes, potentially enabling earlier intervention to prevent subsequent development or exacerbation of balance impairments. Quantifying the stability of the N1 over time could determine how frequently it should be monitored for any potential clinical application, and how long a single measurement could relate to prognosis and inform clinical decision-making. Reliability on short time scales (i.e., days to weeks) would be meaningful for evaluating functional status and determining the initial efficacy of interventions. If the N1 is stable in the absence of an intervention, a change in the N1 following an intervention could offer insight into a specific mechanism mediating a treatment response. If the N1 is relatively stable over a year, it could also be suitable for tracking the longitudinal trajectory of change that is implied by smaller N1 amplitudes in older compared to younger adults (Duckrow, 1999).

One practical consideration for measuring an evoked brain response is the number of trials that are needed to obtain a reliable measurement. Evoked brain responses are typically assessed from averages across trials that are assumed to contain an identical evoked response contaminated by noise and non-time locked neural activity. Averaging across trials preserves the evoked response while reducing the noise that would otherwise confound the measurement of the N1. In theory, more trials will always result in less noise and therefore a more precise estimate of the N1 response. However, in practice, there are diminishing returns to additional trials (Fischer et al., 2017; Meyer et al., 2013; Riesel et al., 2013), which limit the number of conditions that can be assessed, and can introduce additional confounds such as fatigue, changes in attention, or habituation of the evoked brain response (Mierau et al., 2015; Mochizuki et al., 2010; Quintern et al., 1985). Further, from a clinical perspective, there are often time constraints and other considerations (e.g., limited strength, poor balance) that necessitate shorter testing sessions. Internal consistency reliability is a simple method of measuring the consistency of responses across trials to estimate the proportion of the final outcome measure that comes from the N1 relative to the noise that may remain after averaging, and can therefore be used to provide empirically supported practical guidelines on the minimum number of trials needed to obtain a reliable estimate of the N1.

The N1 is often quantified using mobile-brain body imaging (MoBI) approaches that rely on high-density EEG and advanced preprocessing (Makeig et al., 2009), but for clinical applications, a biomarker should be accessible, quickly measured, and easily interpreted. While MoBI approaches were developed to handle the increased frequency and variety of artifacts present during whole-body movement, such as running, it is unclear whether a MoBI approach is necessary to reliably measure the N1. The time required for extensive electrode cap setup (e.g., ≥ 64 channels) and complexity of data cleaning approaches that have yet to be standardized or automated pose significant barriers to clinical translation. While some studies quantify the N1 with a minimal approach using a limited electrode set and simple preprocessing (Duckrow, 1999; Payne et al., 2024, 2023) that could be clinically feasible, no study to date has compared this to a full MoBI approach and whether their reliabilities differ.

Here, we characterized the test-retest reliability and the internal consistency reliability of the N1 elicited by standing balance perturbations in younger and older adults over a one-week interval. A subset of older adults also completed an initial testing session approximately one year prior to enable comparison between short-versus long-term test-retest reliability of the N1. We also examined whether a minimum approach could reliably assess the N1 by comparing two approaches that differ in the number of electrodes and complexity of data preprocessing. These sessions also involved behavioral assessments that have previously been linked to N1 responses (Payne et al., 2022; Payne and Ting, 2020). As an exploratory analysis, we examined the test-retest reliability of behavioral assessments and whether they are sensitive to practice effects.

## 2. Methods

### 2.1 Participants and Experimental Design

10 younger adults (YA) (24 ± 2 years [mean ± standard deviation], 6 identified as female) and 14 older adults (OA) (70 ± 5 years, 6 identified as female) participated in the main experiment involving two testing sessions separated by an interval of 1-7 days (YA: 3.9 ± 2.6 days; OA: 3.5 ± 2.5 days) at the same time of day to control for potential circadian and lifestyle influences. A subset of older adults (n=12, age: 71 ± 5 years, 5 identified as female) also completed one testing session that took place ∼1 year prior to the main experiment (1.18 ± 0.51 years) (Figure 1A). One of these older adults withdrew after the first session of the main experiment due to fatigue and soreness. A second older adult had a longer delay between the two sessions (10 days) due to a fall that happened after the first session. Older adults were included if they were over age 60, had cognitive ability to provide informed consent, could stand without an assistive device for 10 minutes, had no recent history (< 6 months) of pain or injury to their back or lower extremities, and had no history of neurological conditions such as stroke or Parkinson’s disease. The experimental protocol was approved by the Emory University Institutional Review Board, and all participants gave written informed consent before entering the study.

**Figure 1.**
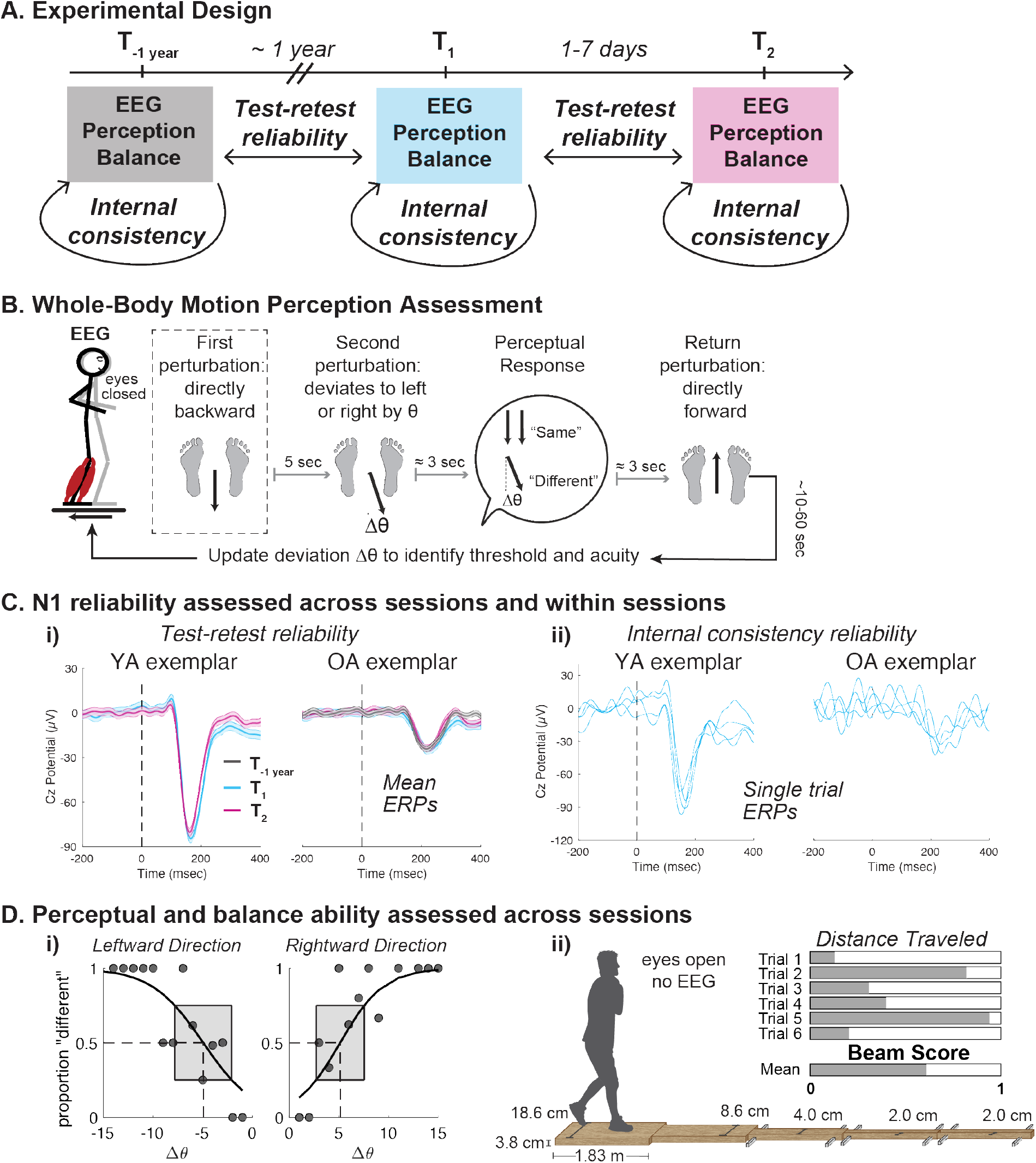
**A**. In the main experiment, younger adults (YA) and older adults (OA) completed two testing sessions within a one-week timespan. A subset of OA also had an initial session ∼1 year prior to the main experiment. B. For each session, participants completed a two-alternative forced choice whole-body motion perception task while EEG was recorded. They were asked to discriminate the direction of a pair of perturbations. The first perturbation was directly backwards, while the second perturbation was backwards with a lateral component (A0) that was adapted online depending on their response. After their verbal response, the platform returned forward. EEG from the first perturbation in each pair was used for analysis (dashed rectangle). C. N1s were compared between sessions using test-retest reliability (i) and within a session using internal consistency reliability (ii). Vertical dashed line denotes perturbation onset. **D**. Perceptual and balance ability were compared between sessions using test-retest reliability, i) Exemplar YA perceptual data for leftward and rightward platform directions. Threshold was defined as the angle at which participants correctly perceived as “different” on half of the trials. Acuity was defined as the range of angles with correct accuracy between 25-75%. ii) Balance ability was quantified as the normalized distance traversed across a narrowing beam, averaged across 6 trials.

### 2.2 Standing balance perturbations

Participants stood barefoot on a platform with their feet hip width apart and arms across their chest. They were exposed to translational support-surface balance perturbations and asked to recover their balance without taking a step. Participants first warmed up with a block of 16-22 perturbations (single ramp-and-hold, 5.1-7.5 cm displacement, 0.8-1 m/s^2^) evenly divided between forward and backward directions with their eyes open, followed by a seated rest break for 10 minutes. They then underwent a two-alternative forced choice perceptual task in which they discriminated whether a pair of perturbations (each 7.5 cm displacement, 15 cm/s velocity, 0.1 m/s^2^ peak acceleration) was in the same or different direction with their eyes closed (Figure 1B) (Bong et al., 2020; Mirdamadi et al., 2024; Puntkattalee et al., 2016). Each trial consisted of a pair of backward perturbations separated by 5 seconds, a verbal perceptual response, and a slow return of the support surface directly forward. The first perturbation occurred at unpredictable timing and was always straight backwards, and the second perturbation deviated slightly from backwards with a lateral component (Δθ). After the second perturbation ended, participants gave a verbal response to indicate whether the two perturbations were in the “same” or “different” direction. Δθ of the second perturbation was adapted online using the psi method implemented through the Palamedes toolbox in MATLAB (Prins and Kingdom, 2018), with leftward and rightward angles interleaved ranging from 0º to 40º in either direction. Up to 60 pairs of perturbations were delivered, with a 5-minute rest break every 15 trials or sooner if requested by the participant. Two older adult participants were unable to complete the full perturbation protocol due to fatigue and only completed 40 pairs of perturbations.

### 2.3 Behavioral assessments

Perceptual ability was assessed in each session using a psychometric function that fit the accuracy of their responses across the range of angles tested. Separate models were performed for leftward and rightward angles. Threshold was defined as the angle at which participants were equally likely to report same or different (Fig 1D, i. – vertical dashed line). Acuity was defined as the interquartile range, or range of angles that spanned 25%-75% accuracy (Fig 1D, i. – shaded rectangle) (Mirdamadi et al., 2024).

Balance ability was assessed using a modified version of the narrowing beam walking test (Sawers and Hafner, 2018). The beam was 9.15 m in length, divided into five fixed-width segments that were each 1.83 m in length. The beam started wide (18.6 cm) and became progressively narrower in discrete intervals (2 cm) (Figure 1D, ii.). At the beginning of the session prior to EEG setup, participants walked across the narrowing beam with their arms across their chest and one foot in front of the other with their eyes open. They started with one foot on the beam and one foot on the floor, and were told to walk as far as they could without losing their balance (i.e., uncrossing their arms or stepping off the beam). Distance traversed was measured as the position of the front of the toes of the last foot on the beam when they uncrossed their arms or stepped off the beam. Balance ability was computed as the average distance traveled normalized to the full beam length. Balance ability was assessed in all sessions for all older adults and for half of the younger adults, but only in one session for the remaining younger adults.

At the end of each session, participants rated their quality of sleep, attention, and fatigue on a Likert scale from 1-10 to determine any transient state-related changes that may influence test-retest reliability.

Participants completed Part A and B of the Trail Making Test as a pen and paper assessment of cognitive set-shifting ability (McKay et al., 2018; Sánchez-Cubillo et al., 2009). Part A involves drawing lines to connect sequential numbers (1-2-3-4), while Part B involves alternating numbers and letters in sequential order (1-A-2-B) without lifting the pen. Set-shifting ability was computed as the difference in completion time between Part B and Part A. Given known practice effects on the Trail Making Test when retested in a short interval (Buck et al., 2008), this was only completed on the second visit of the main experiment. OAs also completed the Trail Making Test one year prior, a sufficient time frame from the main experiment to minimize any practice effects (Basso et al., 1999).

### 2.4 EEG acquisition and analyses

EEG was continuously recorded during standing balance perturbations (sampling frequency 1000Hz, impedance <25kOhm) using a 64-channel active electrode cap (actiCAP) and ActiCHamp amplifier with a 24-bit A/D converter and an online 20 kHz anti-aliasing low-pass filter (Brain Products, GmbH). Fz was the online reference electrode and Fpz was the ground electrode.

EEG analyses were performed in EEGLAB and based on the BeMoBIL Pipeline, a standardized toolbox that is optimized for mobile brain/body imaging data (Klug et al., 2022). Continuous data from the initial practice block and perception block were merged, then downsampled to 250 Hz. BeMoBIL preprocessing consisted of line-noise removal using the zapline plus plugin (Klug and Kloosterman, 2022) and bad channel detection and removal using the cleanrawdata plugin. Since the cleanrawdata plugin uses a random initialization, it was run 20 times and for each iteration identified channels that had correlation with nearby channels < 0.7. A channel was removed if it was flagged as “bad” on more than 7/20 iterations. Bad channels were interpolated and then full-rank average referenced that recovered the online reference (Fz). Preprocessed data were then high-pass filtered, as required for Adaptive mixture component analysis algorithm (AMICA) used to identify maximally-independent components (ICs) (Delorme et al., 2012). Given the discrete nature of the perturbations and variable rest breaks in between trials for participants, AMICA was run on epoched data from -2 to 2 seconds around each perturbation. Since AMICA performance improves with larger amounts of data, we included epochs from the practice blocks, and both the first and second perturbation of each trial pair in the main testing block. However, only perturbations from the first perturbation in each trial pair were used in statistical analyses. ICs were fit to a standard head-model using MNI template, with equivalent current dipoles estimated using the DIPFIT plugin (Oostenveld and Oostendorp, 2002). ICs were removed using the following criteria: any ICs with high residual variance (>20%), ICs that were identified by ICLabel as “eye”, “heart”, or “channel noise”, ICs in which the linear slope of the power spectrum between 2-40 Hz was > -0.33, as commonly seen in muscle artifacts (Fitzgibbon et al., 2016)and any additional ICs labeled as “bad channels” using the TESA toolbox (Rogasch et al., 2017). Finally, powpowcat was used to visually inspect the remaining ICs, to further distinguish brain and muscle ICs (Liu et al., 2024; Studnicki and Ferris, 2023; Thammasan and Miyakoshi, 2020). The resulting solution was applied to the BeMoBIL preprocessed but unfiltered, continuous data in channel space. In the final stage of preprocessing (Figure 2, “Shared preprocessing”), continuous data were referenced to the mastoids, high-pass filtered with a cutoff of 0.5 Hz, low pass filtered with a cutoff of 20 Hz, epoched -2 to 2 seconds, with baseline subtraction -150 to -50 msec (Payne et al., 2019a).

**Figure 2.**
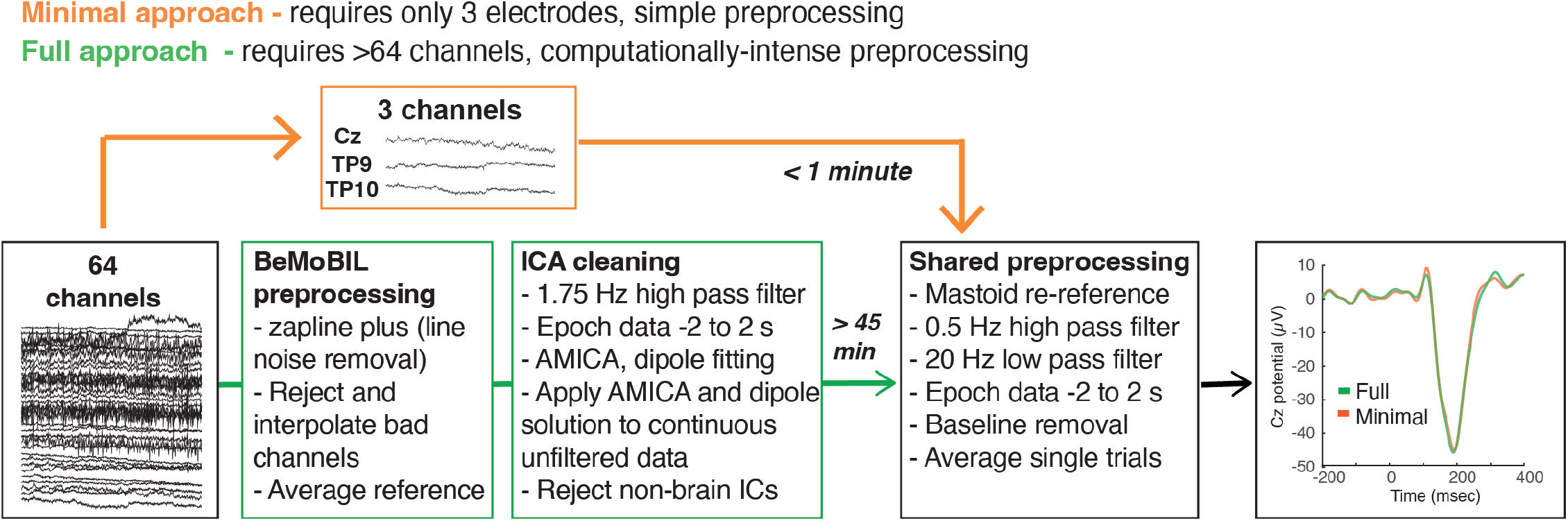
Schematic of different approaches for analyzing N1 amplitude and N1 latency. Despite differences in the number of electrodes and preprocessing complexity and time required, the mean ERP trace generated from each approach was largely overlapping

Prior work has shown that N1s at the Cz electrode have excellent internal consistency using a reduced channel montage and a simple preprocessing pipeline (without AMICA) (Payne et al., 2023). In order to replicate these findings in two independent cohorts, we also characterized the N1 using a minimal approach that was performed with only 3 electrodes (Cz and two mastoids) and simple preprocessing, skipping steps involving the BeMoBIL pipeline and ICA (Figure 2, orange bypass). Preprocessing was identical to that of the final stage of the full approach (“Shared preprocessing”).

We identified the peak N1 amplitude (largest peak negative response) and latency (time of peak negative response) based on the most negative local minimum between 120-200 msec in YA, or 120-300 msec in OA. between 100-300 msec post-perturbation. While most N1s can be identified within 200 msec (Mirdamadi et al., 2024; Payne and Ting, 2020), we extended the upper time bound to 300 msec for older adults since their N1s can be delayed (Duckrow, 1999; Palmer et al., 2024; Payne et al., 2023). For ERPs without a local minimum, the most negative value in the time window was used. Outliers were identified and removed for any trials that exceeded a threshold above or below 3 standard deviations of the mean ERP for 10 consecutive data points between -150 to 300 msec. Across all participants, 3.5% of all trials were removed.

### 2.5 Statistical analyses

N1s were first analyzed using the full approach (Figure 2, green arrows), where amplitude and latency were extracted from the average of all single-trial epochs for each participant and session. To test for group differences in the N1 across sessions, we ran a linear mixed effects model with factors of Group (YA, OA) and Session (T_1_, T_2_). Separate models were run for N1 amplitude and latency.

Test-retest reliability of N1 amplitude and latency were measured using the intraclass correlation coefficient (ICC) estimated with a 2-way mixed effects model with absolute agreement (Shrout and Fleiss, 1979). Separate analyses were performed for each group and testing interval (one week – T_1_ vs T_2_; one year – T_1_ vs T_-1 year_).

Internal consistency was assessed with split-half reliability (Payne et al., 2023). Specifically, N1 amplitudes and latencies were measured from waveforms created by averaging across even and odd subsets of trials within each session for each individual. The consistency of these measures across the even and odd trial subsets was measured (within each group and session) using Pearson’s correlation coefficient corrected by the Spearman-Brown prophecy formula.

Because the full approach is time consuming, requires large amounts of data, and requires expertise that may not be available in a clinical setting (Figure 2, green arrows), we tested whether N1s can be reliably estimated using a minimal approach (Figure 2, orange arrows). We compared N1 amplitudes and latencies obtained using a full approach (64 channels, advanced preprocessing) compared to a minimal approach (3 channels, simple preprocessing) using Pearson correlations. Separate correlations were run for each group and session. All reliability measurements assessed with the full approach were repeated with the minimal approach.

We additionally assessed test-retest reliabilities and internal consistencies as a function of the number of trials included in analyses to provide empirically supported guidelines for determining the number of trials necessary to obtain a reliable measurement of the N1. Analyses were performed iteratively using limited subsets of initial trials (i.e., using only the first 2, 4, 6, etc. trials in the average ERP) to assess the minimum number of trials needed to obtain a reliable estimate of N1 between testing sessions (test-retest reliability) and within testing sessions (internal consistency reliability) (Larson et al., 2010; Meyer et al., 2013). These experiments were designed to collect 60 trials in all individuals, but some individuals were unable to complete the full series due to fatigue. Rather than excluding individuals from these analyses beyond the number of trials they were able to complete, which may overestimate reliabilities, we chose to carry the measurements from the largest number of trials available for those individuals through the analyses up to 60 trials for a more conservative estimate of reliabilities that may be obtained from experiments designed to collect up to 60 trials. These analyses can be used to provide precise guidelines on the minimum number of trials needed for a reliable estimate of the N1 within any experimental condition.

As an exploratory analysis, we also assessed the test-retest reliability of balance ability (normalized beam walking distance) and perceptual ability (threshold and acuity). To examine their sensitivity to practice effects across testing sessions, we ran paired t-tests comparing performance at T_1_ compared to T_2_, and between T_1_ compared to T_-1 year_.

We used commonly reported cutoffs for the reliability coefficients (Portney & Watkins, 2009; Koo & Lee 2016); values < 0.5 indicate poor reliability, values 0.5-0.75 indicate moderate reliability, values 0.75-0.9 indicate good reliability, and values > 0.9 indicate excellent reliability. Analyses were performed using MATLAB and R. Values are reported as mean ± SD unless otherwise stated.

## 3. Results

### 3.1 N1s are smaller and slower in older adults compared to younger adults

Regardless of session, N1s were smaller in older (26.4 ± 9.8 µV) compared to younger adults (44.2 ± 19.3 µV) (p<0.001). Further, N1s were later in older adults (206.6 ± 35.3 ms) compared to younger adults (159 ± 14.4 ms) (p<0.001) (Figure 3i – group ERPs). Each group had high variability between individuals, evidenced by wide shaded regions in group ERP waveforms, for N1 amplitude (younger: 17.5 – 82.0 µV; older: 12.0 – 44.0 µV) and latency (younger: 142 – 186 ms, older: 162 – 282 ms).

**Figure 3.**
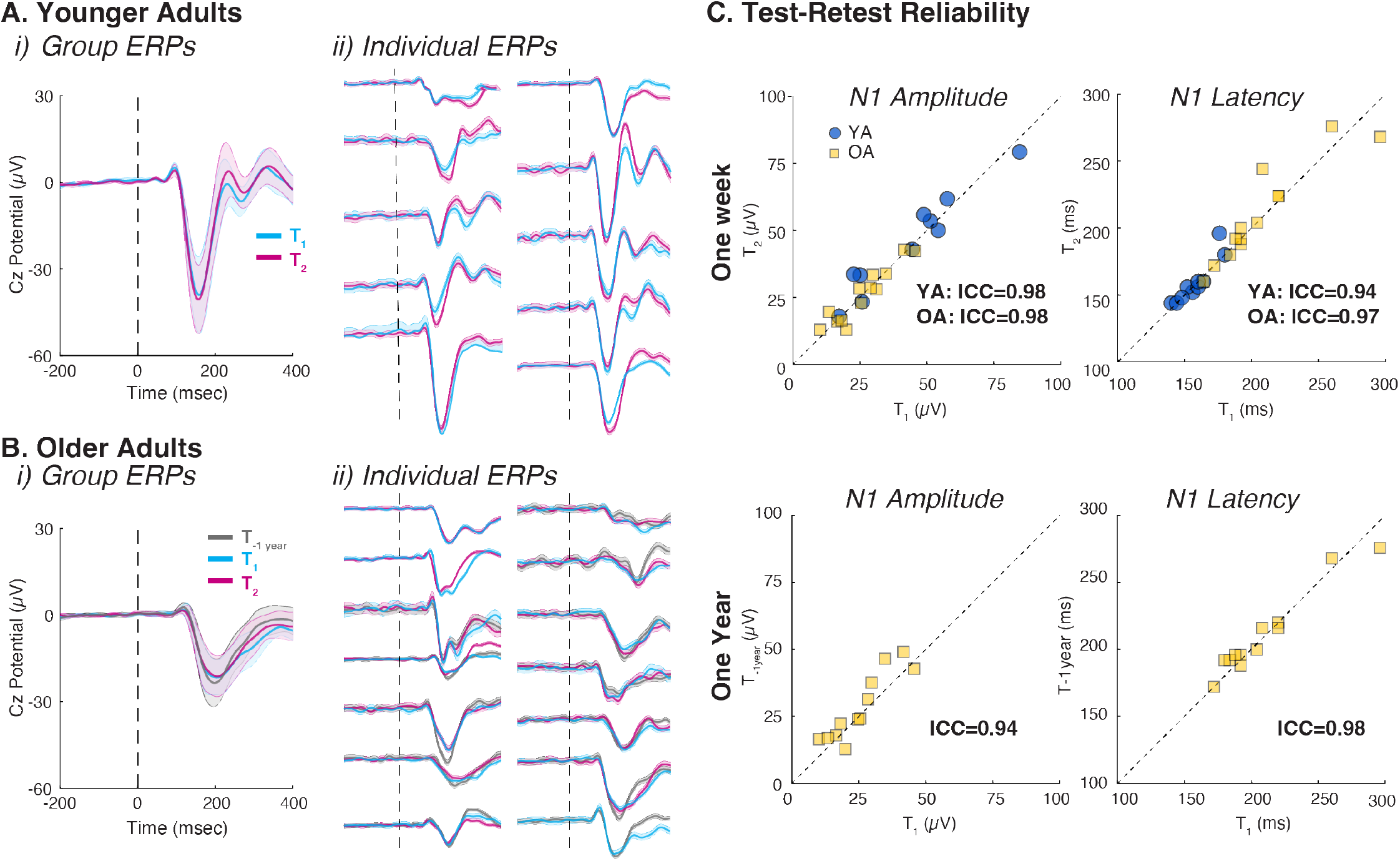
Test-retest reliability across sessions. Perturbation-evoked potentials for YA (A) and OA (B) at the group level (i) and individual level (ii). T_1_ and T_2_ separated by -one week. T_1_ and T_year_ separated by -one year. Solid line represents the mean derived from all trials, and shaded region represents the standard deviation. Vertical dashed line denotes perturbation onset. C. Test-retest reliability for N1 amplitude (left column) and latency (right column) across a one-week (top row) and one-year time span (bottom row). Dashed diagonal denotes line of equality. Full ICC results presented in Table 1.

### 3.2 N1 amplitude and latency have excellent test-retest reliability in YA and OA

Perturbation-evoked potentials varied across individuals but reflected stable cortical activity within an individual across a one-week timespan in younger and older adults (Figure 3 – overlapping blue versus pink traces) and across a one-year timespan in older adults (Figure 3B – overlapping gray traces). ICC analyses confirmed excellent test-retest reliability (ICC>0.9) for N1 amplitude and latency across a one-week and one-year time span (Figure 3 - YA – circles; OA – squares; dashed diagonal denotes line of equality (y=x)). Summary statistics including the ICC and corresponding 95% confidence interval (CI), which were computed separately for each group and retest interval, are summarized in Table 1.

**Table 1.**
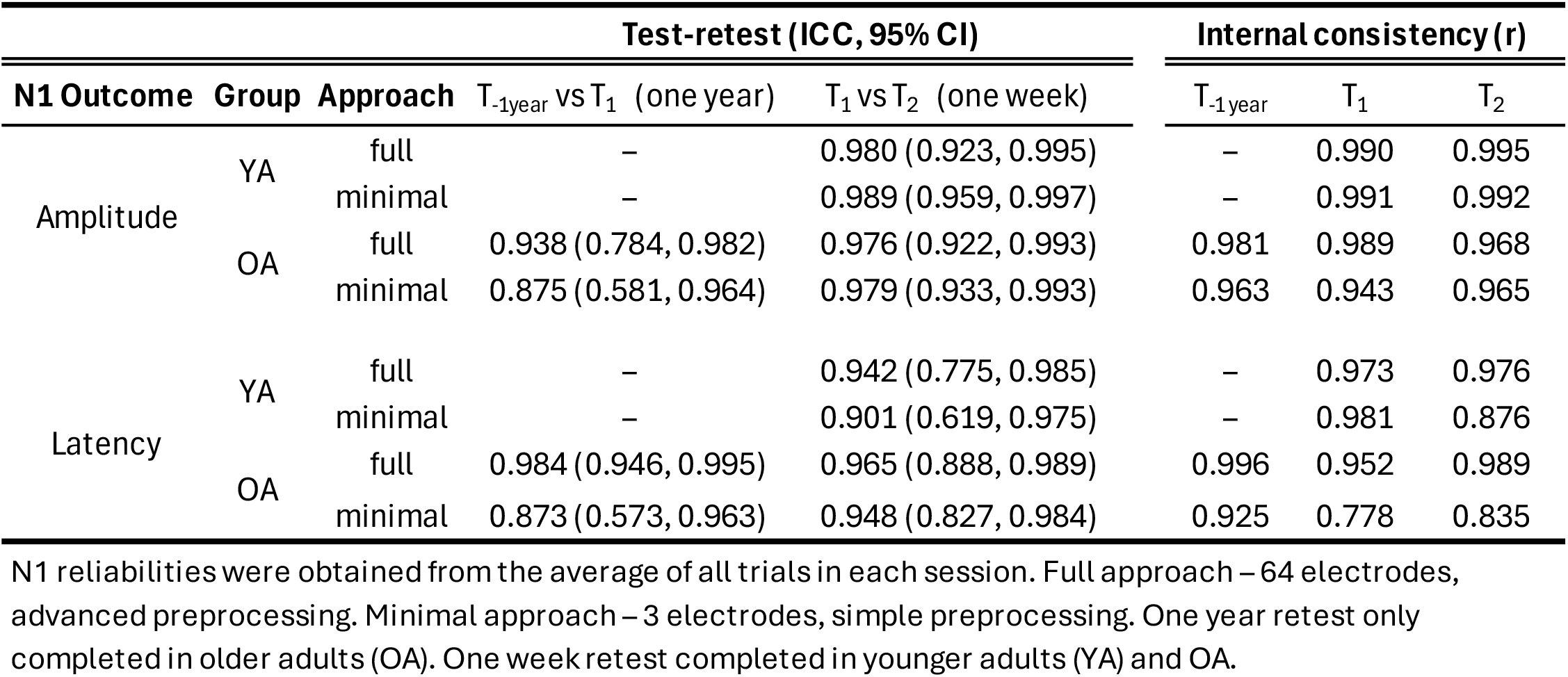
Test-retest reliabilities and internal consistencies for N1 amplitude and latency.

### 3.3 N1 amplitude and latency have excellent internal consistency reliability in YA and OA

Perturbation-evoked potentials appear similar across individual trials (Figure 1C, exemplar YA and exemplar OA). Split-half reliability comparing even vs. odd trials was excellent (r>0.9) for both N1 amplitude and latency for both groups and all sessions (Table 1).

### 3.4 N1s obtained with a minimal approach are similar to those obtained with the full approach

N1 amplitude and latency extracted from the the full and minimal approaches were similar for each session and group (Figure 4A – exemplar participants with overlapping ERP waveforms for the minimal (orange) and full (green) approach. Figure 4B – all participants. YA-circles, OA-squares; gray-T_-1 year_, blue – T_1_, pink – T_2_). Indiviudals with larger N1 amplitudes using the minimal approach had larger N1 amplitudes using the full approach (r= 0.98, p< 0.001; Figure 4B, left). Similarly, indiviudals with slower N1 responses using the minimal approach had slower N1 responses obtained with the full approach (r= 0.93, p<0.001; figure 4B, right). Correlations comparing N1 amplitude and latency between the different approaches for each session and group were all significant (p<0.002; Table 2).

**Table 2.** Pearson correlations comparing N1s obtained using the minimal versus full approach.

|  | Younger Adults |  | Older Adults |  |  |
| --- | --- | --- | --- | --- | --- |
|  | T<sub>1</sub> | T<sub>2</sub> | T<sub>-1year</sub> | T<sub>1</sub> | T<sub>2</sub> |
| **N1 amplitude** | $r=0.994, p<0.001$ | $r=0.960, p<0.001$ | $r=0.946, p<0.001$ | $r=0.962, p<0.001$ | $r=0.981, p<0.001$ |
| **N1 latency** | $r=0.931, p<0.001$ | $r=0.981, p<0.001$ | $r=0.978, p<0.001$ | $r=0.857, p<0.001$ | $r=0.890, p<0.001$ |

**Figure 4.**
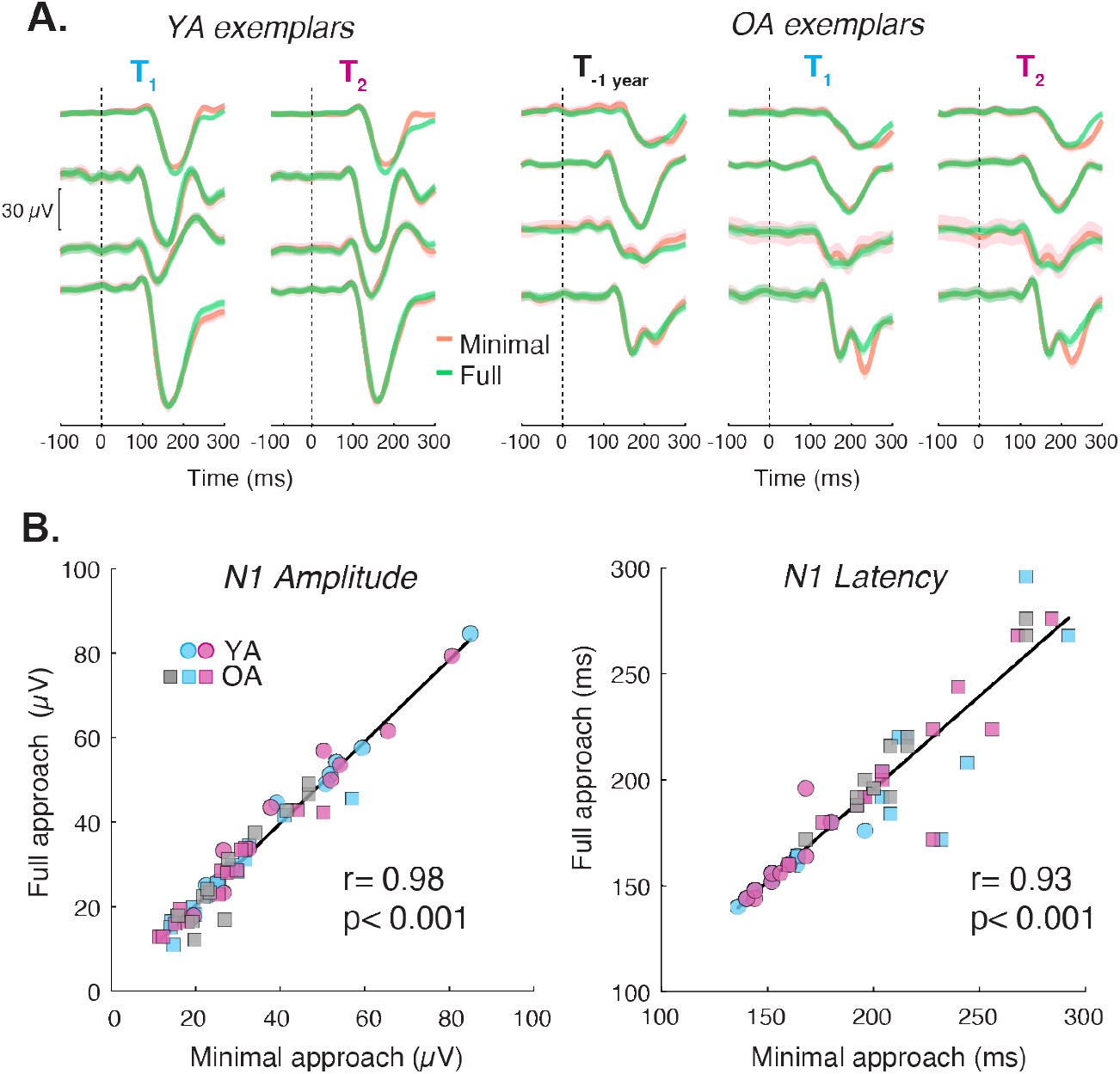
Comparison between N1s derived from the full approach (64 electrodes, advanced preprocessing) versus minimal approach (3 electrodes, simple preprocessing). A. ERPs from the minimal approach (orange) compared to the advanced approach (green) for exemplar YAs and exemplar OAs are highly overlapping. Each row within a group corresponds to ERPs from the same individual on different sessions. B. N1 amplitude (left) and latency (right) using the minimal approach were correlated with those from the advanced approach.

### 3.5 Excellent test-retest reliability and internal consistency were found for N1 amplitude in relatively few trials for both the minimal and full approach

Excellent test-retest reliability (ICC>0.9, Figure 5B left) and internal consistency (r>0.9, Figure 5C left) was generally achieved for N1 amplitude within 6 trials for YA and OA for both the minimal and full approaches. However, more trials were generally required to achieve between good-to-excellent reliabilities for N1 latencies (Figure 5B, C - right), particularly in older adults, as different peaks could be identified across trial subsets within individuals (Figure 5A, OA exemplar bottom row).

**Figure 5.**
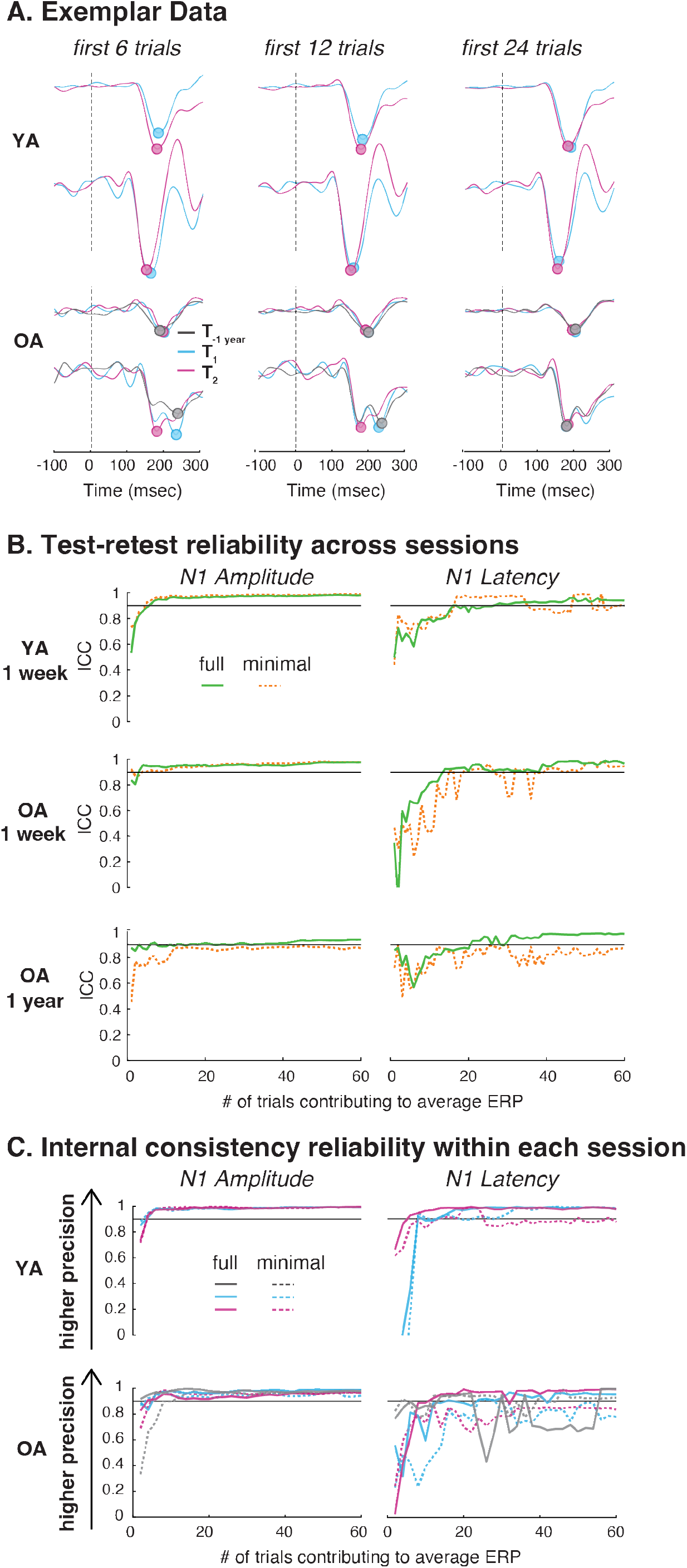
Reliabilities as a function of trial number and approach. A. Exemplar ERPs for younger adults (YA) and older adults (OAs) obtained after averaging across the first 6 trials, 12 trials, or 24 trials for each session. Circles denote peak N1 response that was used to quantify N1 amplitude and latency. YA exemplars and one OA exemplar had one prominent negative peak that was consistent across trials. Another OA exemplar had two different negative peaks which gave rise to different latencies depending upon the number of trials used in the average ERP (e.g., first peak for T_ and second peak for T, and T. _“2“1-1 year_ with ERPs averaged from 6 trials compared to first negative peak at all time points with ERPs averaged from 24 trials). Vertical dashed lines denote perturbation onset. B. Test-retest reliability across one week and one year for N1 amplitude and latency with increasing trial numbers for both the minimal (dashed line) and full approaches (solid line). Black horizontal line denotes threshold for excellent test-retest reliability (ICC>0.9). C. Internal consistency obtained using split-half reliability with increasing trial numbers for N1 amplitude and latency comparing the full approach (solid line) versus minimal approach (dashed line) for each session. Black horizontal line denotes threshold for excellent internal consistency reliability (r>0.9).

### 3.6 Perceptual and balance ability have moderate to excellent test-retest reliability in YA and OA but may be sensitive to practice effects

Test-retest reliabilities for behavioral outcomes (Table 3) were generally lower than those of N1 amplitude and latency (Table 1). Since only half of the YA group had a retest for the beam walking test, we only calculated ICC in OAs. Balance ability had good test-retest reliability for both retest intervals (Figure 6, left column). As shown previously (Mirdamadi et al., 2024), perceptual thresholds and acuity varied across individuals but did not differ between leftward and rightward directions for either session (all p>0.38), and were therefore averaged prior to ICC analysis. Perceptual threshold in both groups and re-test intervals had good test-retest reliability (Figure 6, middle column). Perceptual acuity had moderate to excellent reliability depending upon group and re-test interval (Figure 6, right column). Regardless of group, beam score, perceptual threshold, and perceptual acuity were better at T_2_ compared to T_1_ (p=0.01, p= 0.04, p= 0.01, respectively). Beam score, perceptual threshold, and perceptual acuity were not different between T_-1year_ and T_1_ (all p>0.13). Full summary statistics, including ICC and associated 95% CI for each group and re-test interval are shown in Table 3.

**Table 3.**
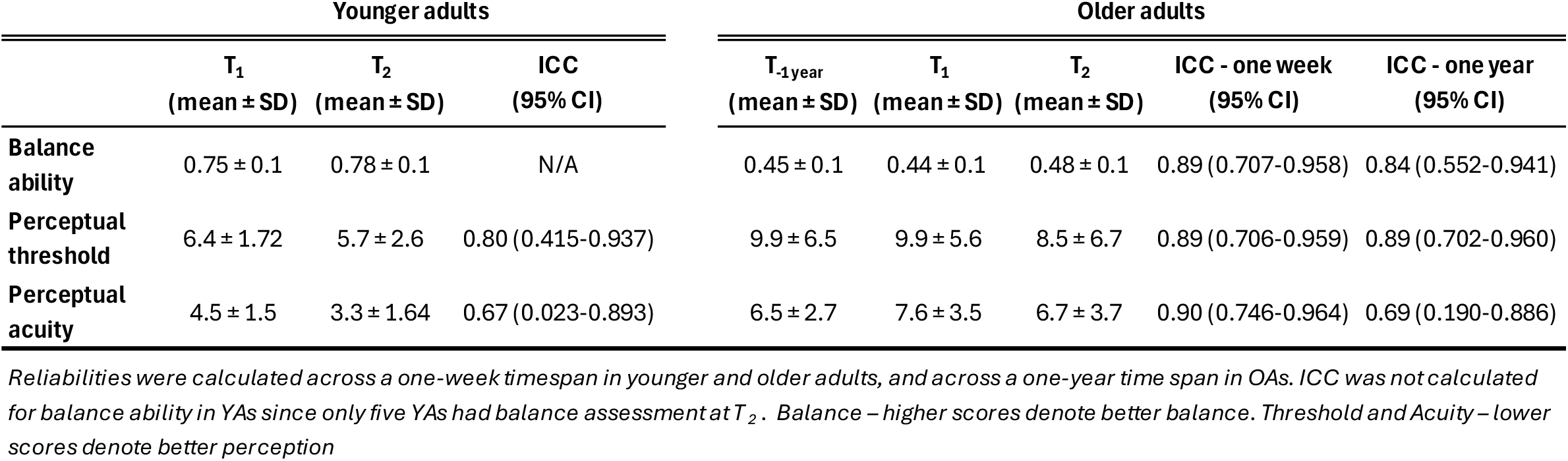
Descriptive results and test-retest reliabilities for balance ability and perceptual ability.

**Figure 6.**
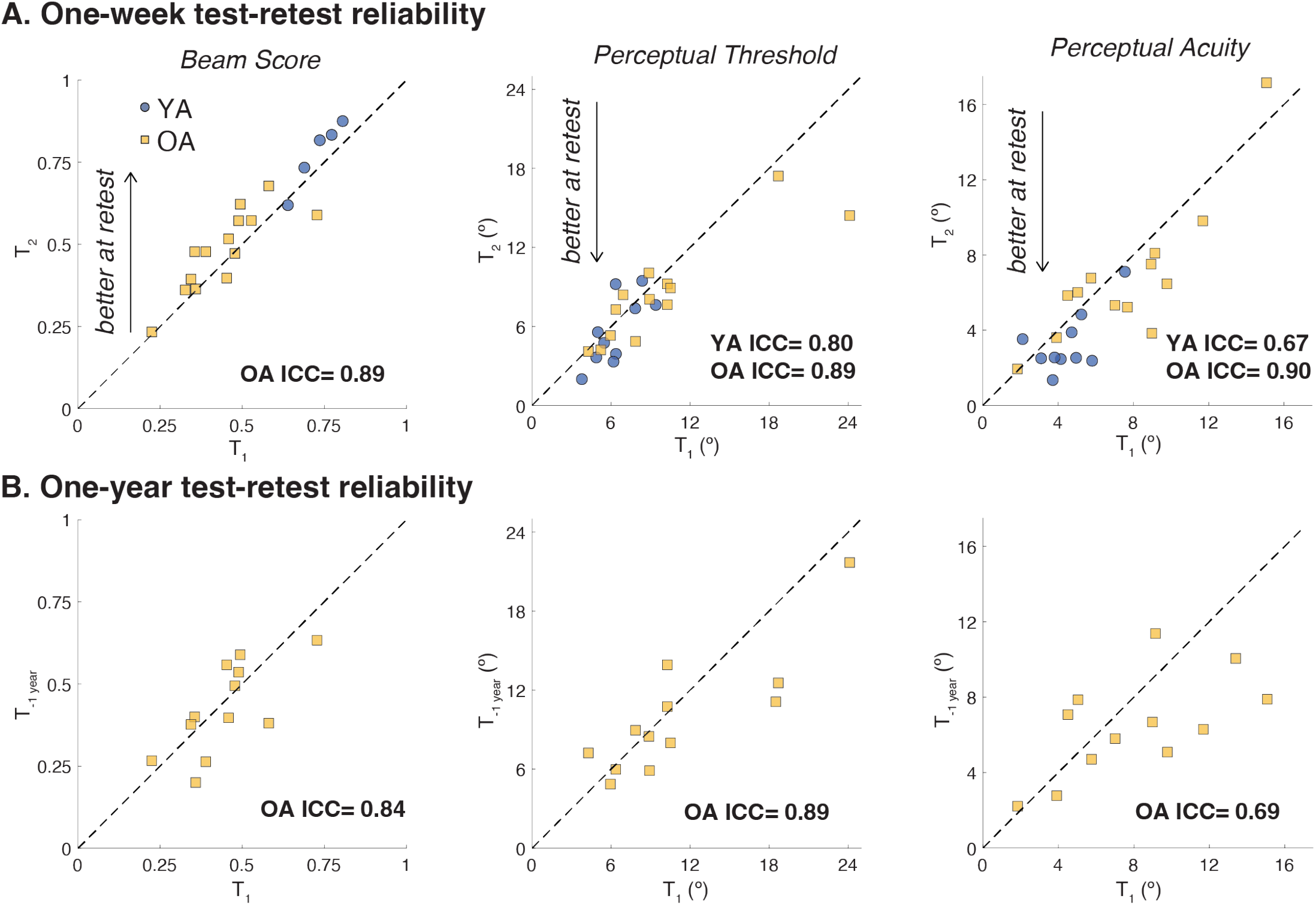
Test-retest reliability for behavioral metrics across a one-week time span **(A)** and a one-year time span (B) (younger adults - blue circles, older adults - yellow squares). Dashed diagonal denotes line of equality. One OA had substantially worse perceptual threshold at T_1_ relative to T_2_, which may have reflected greater self-report of fatigue and soreness after T_r_ Complete ICC and 95% Cl are presented in Table 3.

## 4. DISCUSSION

Perturbation-evoked N1 potentials obtained during a behaviorally-relevant reactive balance task reflect stable individual differences in younger and older adults that can be reliably estimated in as few as six trials. This is the first study to assess test-retest reliability of the N1 response to a balance disturbance, finding individual differences are stable over intervals as long as one year, suggesting the N1 may be suitable for longitudinal investigations of an individual’s functional status. Results also provide precise empirical guidelines for determining the number of trials necessary to obtain a reliable estimate of the N1, and the extent to which that estimate improves with additional trials. We further demonstrate the clinical feasibility of a quick and simple measurement of the N1 by showing that a minimal approach (3 electrodes, simplified preprocessing) results in nearly identical outcome measures (r>0.93) to computationally intense and time-consuming analyses that are standard in MoBI. The excellent stability of the N1 within and between sessions support its usage in individual differences research for determining its potential as a biomarker of balance health and for probing mechanisms underlying interventions.

Individual differences in N1 amplitude and latency were stable across time scales in younger and older adults, reflecting trait-specific cortical activity that could serve as a biomarker of balance health (Olvet and Hajcak, 2008). The later and smaller N1s in older adults compared to younger adults are stable within an individual over a one-year time span, demonstrating its potential use in longitudinal studies tracking age-related or disease-related changes in behavior. Our findings strengthen the interpretation of prior brain-behavior associations that linked individual differences in the N1 to balance ability, balance confidence, and set-shifting ability (Payne et al., 2022, 2021; Payne and Ting, 2020), metrics that have been implicated in fall risk (McKay et al., 2018). Obtaining a stable marker of brain activity that can be mapped to behavior has important implications for guiding clinical decision making. For instance, if changes in brain activity were to precede behavioral declines, the N1 could be used as a monitoring tool to recommend earlier interventions to prevent behavioral decline. Further, since the N1 is stable over time in the absence of any intervention, any changes in the N1 after an intervention could indicate a reliable change in a targeted mechanism rather than random fluctuations or a noisy measurement.

The N1 is known to be modulated by state-related factors such as arousal and fear (Adkin et al., 2008; Sibley et al., 2010), so at first glance the excellent test-retest reliability could be surprising. However, within an individual, the sessions were scheduled at the same time of day to minimize any potential effects of circadian rhythm (Wilson et al., 2014), participants were asked to maintain their normal level of caffeine intake, and the experimental environment was kept as similar as possible across sessions. Although some adults reported large changes in quality of sleep (± 6 out of 10 Likert scale) and hours of sleep (± 4) between sessions, their N1s remained stable, suggesting that any factors that are outside of experimental control and could affect the N1 are likely small relative to the differences in N1 between individuals.

Unlike balance and perception behavior, the N1 was less susceptible to practice effects across sessions, and therefore may represent a more precise and stable index of an individual’s balance health. Nearly all younger and older adults improved their balance and perceptual performance over a one-week timespan, changes that are likely a reflection of task exposure rather than a true change that meaningfully represents the current state of their balance health. Such practice effects are frequently reported on clinical metrics (Buck et al., 2008; Holm et al., 2022; Richardson et al., 2019),and complicate interpretations related to the efficacy or effectiveness of an intervention. In contrast, since the N1 was highly consistent regardless of task exposure with repeated trials or between testing sessions, the N1 has the potential to be a more objective probe of their neurophysiological function for assessing an intervention’s effectiveness at a mechanistic level that could then serve to guide future clinical decision making. Besides practice effects, such behavioral metrics may be sensitive to state-related changes that could vary across sessions. For instance, one older adult reported pain and fatigue in one session but not the other, and presented with a substantial increase in perceptual threshold (nearly double) in that session despite nearly identical N1s. Lastly, some behavioral tests may have floor or ceiling effects and/or little variability between individuals, limiting their potential test-retest reliability. Here, perceptual acuity in the majority of younger adults clustered between 2-5 degrees, likely contributing to the lower ICC for acuity.

Our findings replicate and extend prior work that found excellent internal reliability for N1 *amplitude* elicited by support-surface translations in younger and older adults (Payne et al., 2023) by demonstrating that excellent reliability can be achieved in relatively fewer trials, for both a minimal approach, similar to prior work (Payne et al., 2023), and a full approach commonly used in mobile brain/body imaging (MoBI) research (Makeig et al., 2009). Most MoBI research utilizes high-density EEG and advanced preprocessing pipelines to deal with the increased frequency and types of artifacts present during whole-body behaviors, such as head movement and muscle activity (Klug et al., 2022; Klug and Gramann, 2021; Richer et al., 2024) Since the head doesn’t move until after the N1 in support-surface translations (Payne et al., 2019a, 2023), and the N1 is tightly linked to power in the theta band (Payne et al., 2019b; Solis-Escalante et al., 2021), standard MoBI approaches may be less critical for quantifying the N1, and could hinder clinical translation. Based upon the correlation coefficients comparing N1s obtained from the full vs minimal approach (0.93-0.98), the largest negative impact of choosing the minimal approach would be a 2-7% reduction in the maximum possible correlation between the N1 and other behaviors of interest, which comes with several potential benefits. The minimal approach requires only 3 electrodes and is not dependent upon the number of trials, while the advanced approach requires a full-electrode cap setup and large amounts of data to increase the quality of blind source separation (e.g., AMICA). The ability to obtain a reliable estimate of the N1 with fewer trials offers flexibility in settings that have time constraints or involve populations who are vulnerable to fatigue, while also reducing potential confounds that may occur with longer data collections such as shifts in attention. Finally, the minimal approach can quantify N1s within minutes without extensive training using a fully automated pipeline, and the full approach requires several minutes to hours, including elements of subjective decisions (e.g, IC labeling of brain versus non-brain). However, there may be scenarios in which the full approach is needed, such as source-based analysis or for analyzing power in frequency bands that are known to overlap with muscle artifact, such as beta and gamma power. Further, it’s possible that the full approach is needed in paradigms that elicit larger head movements and/or head movements that occur before the N1. To what extent a minimal approach can be used for other brain metrics or other paradigms that elicit an N1 will require further investigation.

Though test-retest and internal reliability were excellent for N1 *latency* across all trials, reliabilities were generally lower and, unlike N1 amplitude, did not uniformly converge with increasing trial numbers, consistent with prior literature examining ERPs in the cognitive domains (e.g., the error related negativity) (Cassidy et al., 2012; Ip et al., 2018; Lopez et al., 2023). The relatively lower, though still good-to-excellent reliability for N1 latency reflects how some individuals, particularly older adults, have multiple negative peaks during the N1 time window, that could result in different peaks being selected depending upon the subset of trials analyzed (e.g., Figure 5A, bottom row), producing large differences in latency despite similar amplitudes. Two consecutive negative peaks separated by < 100 msec has previously been reported in older adults (Duckrow, 1999), but there is no consensus on the best approach for analysis. Here we chose the most negative peak, even if there was a smaller preceding peak, but it is possible that another method of peak selection could result in more reliable latencies, such as taking the average of the two largest negative peaks. Another method is manual peak detection, but this comes at a tradeoff of inter-rater differences due to observer biases, time required for training and visual inspection, and reproducibility (Delorme, 2023). In the context of developing a biomarker and enhancing replication, we believe that having a robust, automated pipeline outweighs potential benefits that could come with manual methods, even if that means collecting additional trials. Whatever approach is used, reliability will be influenced by how N1s are defined. Given large inter-individual variability in ERP shape, we recommend that future research clearly define their method of peak selection, including the time window and how multiple negative peaks in the time window are handled. Source-based analyses may offer insight into whether multiple negative peaks in the N1 time window are originating from different sources, which will not only advance our understanding of the N1, but also help refine approaches to determine its reliability.

While N1 amplitude achieved excellent reliability within 6 trials, it’s important to consider that reliability is context-dependent and will need to be quantified on a study-by-study basis (Clayson and Miller, 2017). Specifically, our results may not generalize to clinical populations, to N1s quantified during other types of tasks, or to N1s elicited by other perturbation methods that may result in more prominent artifacts or head movements. Unlike test-retest reliability which requires repeated experimental sessions, internal reliability can be quantified and reported in any EEG study. Because reliability limits statistical power, a metric with poor reliability could preclude or falsely magnify group- or task-related differences in brain activity or brain-behavior correlations (Clayson, 2024; Clayson and Miller, 2017; Hajcak et al., 2017). Routine reporting of internal consistency is needed to enhance reproducibility, interpretation of results, and optimize future experimental designs. The minimum number of trials needed for excellent reliability found here is not a hard rule, but is rather meant to serve as a reference for future studies and will need to be evaluated on a study-by-study basis. For instance, if N1 latency is a desired outcome, we suggest that researchers over-estimate the minimum number of trials, based on our current results and prior literature (Lopez et al., 2023). However, if the primary outcome of a study is N1 amplitude, understanding the diminishing returns of additional trials could motivate the inclusion of more experimental conditions rather than excessive repetition of trials within any condition.

Another important consideration in determining reliabilities as a function of trial number is the role of practice and familiarization of the task. Some studies have suggested to remove the first trial (or first few trials) to control for transient emotional changes (e.g., anxiety, arousal) that occur with the novelty of a task (Maki and Whitelaw, 1993; Zaback et al., 2021). One limitation of our study is that we had a practice block of perturbations that differed in several factors from perturbations in the main experiment, so it is unclear if and to what extent practice effects could have impacted N1 reliability. However, the practice block was intentional to allow participants to adapt to the perturbations, prior to adding complexity with a perception task used in the main experiment. For studies in which practice trials are similar to their main paradigm, it would be helpful to quantify any potential practice effects to justify whether initial trials are included or excluded to strengthen the interpretation of their main results of interest. Future research specifically designed to test the role of practice trials is needed to understand their impact on the N1.

### Conclusion and future directions

The N1 is stable across a timespan of one week and one year, can be reliably estimated with relatively few trials using a minimal approach, and can be quickly quantified within minutes using entirely automated procedures, each of which is critical for translating any biomarker into clinical practice. The current results provide empirical guidance for reliably measuring the balance N1 elicited by support-surface translations. However, as reliability may differ across tasks, settings, and populations, we recommend reporting internal consistencies of N1s in all studies to inform the design of more powerful investigations of the N1. Since N1s are stable in the absence of interventions over a one-year time span in older adults, it may be possible to develop longitudinal trajectories of the N1 to detect changes in functional status prior to the onset of behavioral declines. As prior cross-sectional studies have linked individual differences in N1 to behaviors that predict falls (McKay et al., 2018; Payne et al., 2022, 2021), our findings demonstrating its excellent reliability lay the foundation to determine its predictive validity, or extent to which the N1 serves as a biomarker for behaviors of interest (e.g., fall risk).

## References

Adkin, A.L., Campbell, A.D., Chua, R., Carpenter, M.G., 2008. The influence of postural threat on the cortical response to unpredictable and predictable postural perturbations. Neurosci Lett 435, 120–125. 10.1016/j.neulet.2008.02.018

Adkin, A.L., Quant, S., Maki, B.E., McIlroy, W.E., 2006. Cortical responses associated with predictable and unpredictable compensatory balance reactions. Experimental Brain Research 172, 85–93. 10.1007/s00221-005-0310-9

Basso, M.R., Bornstein, R.A., Lang, J.M., 1999. Practice effects on commonly used measures of executive function across twelve months. Clin Neuropsychol 13, 283–292. 10.1076/clin.13.3.283.1743

Beason-Held, L.L., Goh, J.O., An, Y., Kraut, M.A., O’Brien, R.J., Ferrucci, L., Resnick, S.M., 2013. Changes in Brain Function Occur Years before the Onset of Cognitive Impairment. J. Neurosci. 33, 18008–18014. 10.1523/JNEUROSCI.1402-13.2013

Berger, W., Quintern, J., Dietz, V., 1987. Afferent and efferent control of stance and gait: developmental changes in children. Electroencephalography and Clinical Neurophysiology 66, 244–252. 10.1016/0013-4694(87)90073-3

Bong, S.M., McKay, J.L., Factor, S.A., Ting, L.H., 2020. Perception of whole-body motion during balance perturbations is impaired in Parkinson’s disease and is associated with balance impairment. Gait & Posture 76, 44–50. 10.1016/j.gaitpost.2019.10.029

Buck, K.K., Atkinson, T.M., Ryan, J.P., 2008. Evidence of practice effects in variants of the Trail Making Test during serial assessment. J Clin Exp Neuropsychol 30, 312–318. 10.1080/13803390701390483

Cassidy, S.M., Robertson, I.H., O’Connell, R.G., 2012. Retest reliability of event-related potentials: Evidence from a variety of paradigms. Psychophysiology 49, 659–664. 10.1111/j.1469-8986.2011.01349.x

Clayson, P.E., 2024. The psychometric upgrade psychophysiology needs. Psychophysiology 61, e14522. 10.1111/psyp.14522

Clayson, P.E., Miller, G.A., 2017. Psychometric considerations in the measurement of event-related brain potentials: Guidelines for measurement and reporting. International Journal of Psychophysiology, Rigor and Replication: Towards Improved Best Practices in Psychophysiological Research 111, 57–67. 10.1016/j.ijpsycho.2016.09.005

Delorme, A., 2023. EEG is better left alone. Sci Rep 13, 2372. 10.1038/s41598-023-27528-0

Delorme, A., Palmer, J., Onton, J., Oostenveld, R., Makeig, S., 2012. Independent EEG Sources Are Dipolar. PLoS One 7. 10.1371/journal.pone.0030135

Duckrow, R., 1999. Stance perturbation-evoked potentials in old people with poor gait and balance. Clinical Neurophysiology 110, 2026–2032. 10.1016/S1388-2457(99)00195-9

Fischer, A.G., Klein, T.A., Ullsperger, M., 2017. Comparing the error-related negativity across groups: The impact of error- and trial-number differences. Psychophysiology 54, 998–1009. 10.1111/psyp.12863

Fitzgibbon, S.P., DeLosAngeles, D., Lewis, T.W., Powers, D.M.W., Grummett, T.S., Whitham, E.M., Ward, L.M., Willoughby, J.O., Pope, K.J., 2016. Automatic determination of EMG-contaminated components and validation of independent component analysis using EEG during pharmacologic paralysis. Clinical Neurophysiology 127, 1781–1793. 10.1016/j.clinph.2015.12.009

Hajcak, G., Meyer, A., Kotov, R., 2017. Psychometrics and the neuroscience of individual differences: Internal consistency limits between-subjects effects. J Abnorm Psychol 126, 823–834. 10.1037/abn0000274

Holm, S.P., Wolfer, A.M., Pointeau, G.H.S., Lipsmeier, F., Lindemann, M., 2022. Practice effects in performance outcome measures in patients living with neurologic disorders – A systematic review. Heliyon 8, e10259. 10.1016/j.heliyon.2022.e10259

Ip, C.-T., Ganz, M., Ozenne, B., Sluth, L.B., Gram, M., Viardot, G., l’Hostis, P., Danjou, P., Knudsen, G.M., Christensen, S.R., 2018. Pre-intervention test-retest reliability of EEG and ERP over four recording intervals. Int J Psychophysiol 134, 30–43. 10.1016/j.ijpsycho.2018.09.007

Klug, M., Gramann, K., 2021. Identifying key factors for improving ICA-based decomposition of EEG data in mobile and stationary experiments. European Journal of Neuroscience 54, 8406–8420. 10.1111/ejn.14992

Klug, M., Jeung, S., Wunderlich, A., Gehrke, L., Protzak, J., Djebbara, Z., Argubi-Wollesen, A., Wollesen, B., Gramann, K., 2022. The BeMoBIL Pipeline for automated analyses of multimodal mobile brain and body imaging data. 10.1101/2022.09.29.510051

Klug, M., Kloosterman, N.A., 2022. Zapline-plus: a Zapline extension for automatic and adaptive removal of frequency-specific noise artifacts in M/EEG. 10.1101/2021.10.18.464805

Larson, M.J., Baldwin, S.A., Good, D.A., Fair, J.E., 2010. Brief Reports: Temporal stability of the error-related negativity (ERN) and post-error positivity (Pe): The role of number of trials. Psychophysiology 47, 1167– 1171. 10.1111/j.1469-8986.2010.01022.x

Liu, C., Downey, R.J., Salminen, J.S., Arvelo Rojas, S., Richer, N., Pliner, E.M., Hwang, J., Cruz-Almeida, Y., Manini, T.M., Hass, C.J., Seidler, R.D., Clark, D.J., Ferris, D.P., 2024. Electrical brain activity during human walking with parametric variations in terrain unevenness and walking speed. Imaging Neuroscience 2, 1–33. 10.1162/imag_a_00097

Lopez, K.L., Monachino, A.D., Vincent, K.M., Peck, F.C., Gabard-Durnam, L.J., 2023. Stability, change, and reliable individual differences in electroencephalography measures: A lifespan perspective on progress and opportunities. NeuroImage 275, 120116. 10.1016/j.neuroimage.2023.120116

Makeig, S., Gramann, K., Jung, T.-P., Sejnowski, T.J., Poizner, H., 2009. Linking brain, mind and behavior. International Journal of Psychophysiology, Neural Processes in Clinical Psychophysiology 73, 95–100. 10.1016/j.ijpsycho.2008.11.008

Maki, B.E., Whitelaw, R.S., 1993. Influence of expectation and arousal on center-of-pressure responses to transient postural perturbations. Journal of Vestibular Research 3, 25–39.

McEvoy, L.K., Smith, M.E., Gevins, A., 2000. Test-retest reliability of cognitive EEG. Clin Neurophysiol 111, 457–463. 10.1016/s1388-2457(99)00258-8

McKay, J.L., Lang, K.C., Ting, L.H., Hackney, M.E., 2018. Impaired set shifting is associated with previous falls in individuals with and without Parkinson’s disease. Gait and Posture. 10.1016/j.gaitpost.2018.02.027

Meyer, A., Riesel, A., Hajcak Proudfit, G., 2013. Reliability of the ERN across multiple tasks as a function of increasing errors. Psychophysiology 50, 1220–1225. 10.1111/psyp.12132

Mierau, A., Hülsdünker, T., Strüder, H.K., 2015. Changes in cortical activity associated with adaptive behavior during repeated balance perturbation of unpredictable timing. Front Behav Neurosci 9. 10.3389/fnbeh.2015.00272

Mirdamadi, J.L., Ting, L.H., Borich, M.R., 2024. Distinct Cortical Correlates of Perception and Motor Function in Balance Control. J Neurosci 44, e1520232024. 10.1523/JNEUROSCI.1520-23.2024

Mochizuki, G., Boe, S., Marlin, A., McIlroy, W.E., 2010. Perturbation-evoked cortical activity reflects both the context and consequence of postural instability. Neuroscience 170, 599–609. 10.1016/j.neuroscience.2010.07.008

Mochizuki, G., Sibley, K.M., Cheung, H.J., Camilleri, J.M., McIlroy, W.E., 2009. Generalizability of perturbation-evoked cortical potentials: Independence from sensory, motor and overall postural state. Neuroscience Letters 451, 40–44. 10.1016/j.neulet.2008.12.020

Olvet, D., Hajcak, G., 2008. The error-related negativity (ERN) and psychopathology: Toward an endophenotype. Clinical Psychology Review 28, 1343–1354. 10.1016/j.cpr.2008.07.003

Oostenveld, R., Oostendorp, T.F., 2002. Validating the boundary element method for forward and inverse EEG computations in the presence of a hole in the skull. Hum Brain Mapp 17, 179–192. 10.1002/hbm.10061

Palmer, J.A., Payne, A.M., Mirdamadi, J.L., Ting, L.H., Borich, M.R., 2024. Delayed Cortical Responses During Reactive Balance After Stroke Associated With Slower Kinetics and Clinical Balance Dysfunction. Neurorehabil Neural Repair 15459683241282786. 10.1177/15459683241282786

Payne, A.M., Hajcak, G., Ting, L.H., 2019a. Dissociation of muscle and cortical response scaling to balance perturbation acceleration. Journal of Neurophysiology 121, 867–880. 10.1152/jn.00237.2018

Payne, A.M., Hajcak, G., Ting, L.H., 2018. Dissociation of muscle and cortical response scaling to balance perturbation acceleration. Journal of Neurophysiology 121, 867–880. 10.1152/jn.00237.2018

Payne, A.M., McKay, J.L., Ting, L.H., 2022. The cortical N1 response to balance perturbation is associated with balance and cognitive function in different ways between older adults with and without Parkinson’s disease. Cerebral Cortex Communications 3, tgac030. 10.1093/texcom/tgac030

Payne, A.M., Palmer, J.A., McKay, J.L., Ting, L.H., 2021. Lower Cognitive Set Shifting Ability Is Associated With Stiffer Balance Recovery Behavior and Larger Perturbation-Evoked Cortical Responses in Older Adults. Front Aging Neurosci 13, 742243. 10.3389/fnagi.2021.742243

Payne, A.M., Schmidt, N.B., Meyer, A., Hajcak, G., 2024. The balance N1 is larger in anxious children and associated with the error-related negativity. Biological Psychiatry Global Open Science 100393. 10.1016/j.bpsgos.2024.100393

Payne, A.M., Ting, L.H., 2020. Worse balance is associated with larger perturbation-evoked cortical responses in healthy young adults. Gait & Posture 80, 324–330. 10.1016/j.gaitpost.2020.06.018

Payne, A.M., Ting, L.H., Hajcak, G., 2023. The balance N1 and the ERN correlate in amplitude across individuals in small samples of younger and older adults. Exp Brain Res 241, 2419–2431. 10.1007/s00221-023-06692-9

Payne, A.M., Ting, L.H., Hajcak, G., 2019b. Do sensorimotor perturbations to standing balance elicit an error-related negativity? Psychophysiology 56, e13359. 10.1111/psyp.13359

Prins, N., Kingdom, F.A.A., 2018. Applying the Model-Comparison Approach to Test Specific Research Hypotheses in Psychophysical Research Using the Palamedes Toolbox. Frontiers in Psychology 9.

Puntkattalee, M.J., Whitmire, C.J., Macklin, A.S., Stanley, G.B., Ting, L.H., 2016. Directional acuity of whole-body perturbations during standing balance. Gait & Posture 48, 77–82. 10.1016/j.gaitpost.2016.04.008

Purohit, R., Bhatt, T., 2022. Mobile Brain Imaging to Examine Task-Related Cortical Correlates of Reactive Balance: A Systematic Review. Brain Sci 12, 1487. 10.3390/brainsci12111487

Quant, S., Adkin, A.L., Staines, W.R., Maki, B.E., McIlroy, W.E., 2004. The effect of a concurrent cognitive task on cortical potentials evoked by unpredictable balance perturbations. BMC Neurosci 5, 18. 10.1186/1471-2202-5-18

Quintern, J., Berger, W., Dietz, V., 1985. Compensatory reactions to gait perturbations in man: Short- and long-term effects of neuronal adaptation. Neuroscience Letters 62, 371–375. 10.1016/0304-3940(85)90577-4

Richardson, E., Burnell, J., Adams, H.R., Bohannon, R.W., Bush, E.N., Campbell, M., Chen, W.H., Coons, S.J., Papadopoulos, E., Reeve, B.R., Rooks, D., Daniel, G., 2019. Developing and Implementing Performance Outcome Assessments: Evidentiary, Methodologic, and Operational Considerations. Drug Inf J 53, 146–153. 10.1177/2168479018772569

Richer, N., Bradford, J.C., Ferris, D.P., 2024. Mobile neuroimaging: What we have learned about the neural control of human walking, with an emphasis on EEG-based research. Neuroscience & Biobehavioral Reviews 162, 105718. 10.1016/j.neubiorev.2024.105718

Riesel, A., Weinberg, A., Endrass, T., Meyer, A., Hajcak, G., 2013. The ERN is the ERN is the ERN? Convergent validity of error-related brain activity across different tasks. Biol Psychol 93, 377–385. 10.1016/j.biopsycho.2013.04.007

Rogasch, N.C., Sullivan, C., Thomson, R.H., Rose, N.S., Bailey, N.W., Fitzgerald, P.B., Farzan, F., Hernandez-Pavon, J.C., 2017. Analysing concurrent transcranial magnetic stimulation and electroencephalographic data: A review and introduction to the open-source TESA software. NeuroImage 147, 934–951. 10.1016/j.neuroimage.2016.10.031

Sánchez-Cubillo, I., Periáñez, J.A., Adrover-Roig, D., Rodríguez-Sánchez, J.M., Ríos-Lago, M., Tirapu, J., Barceló, F., 2009. Construct validity of the Trail Making Test: role of task-switching, working memory, inhibition/interference control, and visuomotor abilities. J Int Neuropsychol Soc 15, 438–450. 10.1017/S1355617709090626

Sawers, A., Hafner, B., 2018. Validation of the Narrowing Beam Walking Test in Lower Limb Prosthesis Users. Arch Phys Med Rehabil 99, 1491-1498.e1. 10.1016/j.apmr.2018.03.012

Shrout, P.E., Fleiss, J.L., 1979. Intraclass correlations: uses in assessing rater reliability. Psychol Bull 86, 420– 428. 10.1037//0033-2909.86.2.420

Sibley, K.M., Mochizuki, G., Frank, J.S., McIlroy, W.E., 2010. The relationship between physiological arousal and cortical and autonomic responses to postural instability. Exp Brain Res 203, 533–540. 10.1007/s00221-010-2257-8

Solis-Escalante, T., Stokkermans, M., Cohen, M.X., Weerdesteyn, V., 2021. Cortical responses to whole-body balance perturbations index perturbation magnitude and predict reactive stepping behavior. Eur J Neurosci 54, 8120–8138. 10.1111/ejn.14972

Studnicki, A., Ferris, D.P., 2023. Parieto-Occipital Electrocortical Dynamics during Real-World Table Tennis. eNeuro 10, ENEURO.0463-22.2023. 10.1523/ENEURO.0463-22.2023

Thammasan, N., Miyakoshi, M., 2020. Cross-Frequency Power-Power Coupling Analysis: A Useful Cross-Frequency Measure to Classify ICA-Decomposed EEG. Sensors (Basel) 20, 7040. 10.3390/s20247040

Varghese, J.P., McIlroy, R.E., Barnett-Cowan, M., 2017. Perturbation-evoked potentials: Significance and application in balance control research. Neuroscience & Biobehavioral Reviews 83, 267–280. 10.1016/j.neubiorev.2017.10.022

Wilson, T.W., Heinrichs-Graham, E., Becker, K.M., 2014. Circadian modulation of motor-related beta oscillatory responses. NeuroImage 102, 531–539. 10.1016/j.neuroimage.2014.08.013

Zaback, M., Reiter, E.R., Adkin, A.L., Carpenter, M.G., 2021. Initial experience of balance assessment introduces ‘first trial’ effects on emotional state and postural control. Gait & Posture 88, 116–121. 10.1016/j.gaitpost.2021.05.013

